# Placement of new comb cells built by honeybees is guided by sub-cell scale features to align with the existing layout

**DOI:** 10.1101/2022.07.13.499858

**Authors:** Vincent Gallo, Alice Bridges, Joseph Woodgate, Lars Chittka

## Abstract

The comb of honeybees has long been the subject of curiosity and admiration. Its noteworthy features include the even hexagonal layout and the sharing of walls, both side walls and bases, that provide a maximum storage volume while using a minimum of wax for its construction. The efficiency of its structure relies on a regular layout where cells are positioned correctly relative to each other. Each new cell should be placed exactly between two previously constructed cells, a task made more difficult by the incomplete nature of cells at the edge of comb where the new ones are to be built. By offering bees shaped wax stimuli we have identified sub-cell scale shapes that trigger certain patterns of behaviour by honeycomb construction workers. These behaviours, we found, cause the creation of a new cell to be aligned to small concavities and for cell walls to be built mid-way between two such stimuli. We show that by responding to the form of partially constructed comb, the bees build the next iteration of cells and walls at the locations necessary for the repeated pattern of cells to continue.

## Introduction

This chapter presents experiments to test predictions concerning the behaviour of worker bees when building the early elements of comb, before the full formation of each cell. Measurements, description and analysis of completed cells that comprise comb are plentiful, eg. (Hepburn et al. 1991; Hepburn and Whiffler 1991; Smith et al. 2021; Yang et al. 2021), but nascent cells are less well-characterised. Observing the process of comb construction is difficult, even when using an observation hive, as both the construction workers and the workpiece are typically covered by other bees. The impossibility of direct observation was noted by Huber, whose solution was to prevent construction either downwards or sideways, forcing comb to be built upwards (Huber 1814:136). By so doing, Huber had arranged for the point of construction to be raised above the mass of bees which allowed him to observe the process. As a result, Huber was able to describe the initial deposition and sculpting of wax that eventually formed two rows of cells.

The first action leading to a cell, as noted by Huber, was the removal of wax to enlarge a small indentation. The following step was the addition of wax to extend the dished area until “…the diameter of the cavity was equal to that of an ordinary cell…” whereupon he noted that wax was added to the periphery. Work progressed at multiple sites, allowing Huber to observe the conjunction of two nascent cells as “two adjacent cavities… separated only by a common edge, formed from the gathering together of the wax particles drawn from their interior” (1814:141).

The initial stages of cell construction have also been modelled using a computer simulation of cell layout, which assumed that a cell base, a shallow dish, would be formed by expansion of an existing inter-cell cleft (Nazzi 2016). Nazzi continued his model sequence by assuming that the cell base, thus formed, would be extended until it was an appropriate size to trigger the construction of a surrounding wall.

The descriptions by Huber, and the model by Nazzi, are both refer to construction at the edge of comb focused on a clef formed where the walls of two extant cells meet. Our explanation of the mechanism that addresses both Huber’s description and Nazzi’s model, is provided by two hypotheses. The outer surface at the point where two cells meet yields a concave site which, according to our first hypothesis, will trigger a reaction by the builders to extend the depression. Furthermore, the edge of the perceived depression, the point at which the existing cell’s walls turn away, will trigger the reaction to deposit wax around the edge and will eventually to become a wall.

We predict that there will be stigmergic reactions to perceived conditions based on the current state of construction. The assumption that cell construction behaviour would proceed according to these hypotheses leads to three predictions, all pertinent to the beginning stages of cell construction, are:

P1 Wax deposition will begin at the edges of a stimulus comprising a shallow depression, leading to the eventual location of cell walls.
P2 Two shallow depressions will both attract wax depositions at their rim, and there will likely be a deposition that leads to a cell wall, at the mid-way point between the two. The resulting wall will lie orthogonal to the line connecting the centre points.
P3 A stimulus in the form of a low wall will initiate the formation of the remaining cell walls around a cell-sized enclosure. Two such stimuli joined at a V-shape will cause two cells to be formed, and both will result in a new wall being built at the cell intersection. The resulting wall will lie at an angle that bisects the stimulus “V”.

To test these predictions, we fashioned stimuli comprising wax forms where each was designed to trigger one of the predicted behaviours. We then placed the stimuli in hives leaving the bees to build honeycomb upon them. These samples were inspected periodically and, when appropriate, the alignment and position of cell walls were measured for comparison with that of the stimuli.

## Methods

### Hive handling and recording

#### Hives

The studies reported here were all conducted during May, June and July 2020 and June and July 2021 at an apiary in Reigate, England (51.23°N, 0.19°W) using colonies of honeybees, *Apis mellifera*. No attempt was made to identify the sub-species as these colonies were all the product of several generations of locally reared queens mated during uncontrolled flights in the area local to the apiary. The colonies were housed in Modified British National hives comprising an open mesh floor and a single brood box housing 11 frames; including 10 conventional frames plus one used to carry the experimental wax stimulus. All hives were configured in ‘warm’ alignment, that is with the frames set transverse to the entrance. The test frame was placed in each hive as the 7^th^ frame from the front, which was, for the period of the experiments, the edge of the brood area.

#### Instigation of comb construction

In England, the conditions were variable during these experiments, with periods of inclement weather. Observations at the hive entrances and inspection of stores suggested that for some of the time there was a plentiful supply of natural nectar, but at others reduced activity suggested that forage was limited. To overcome the lean periods the hives were also equipped with *ad libitum* 1:1 sucrose solution (1.0 kg cane sugar in 1.0 l water). With both supplementary feed and natural forage available, the colonies were inclined to extend their storage space by building honeycomb.

#### Wax

The wax used for both the vertical backplane and any detailed adornment was from the hives managed within the same apiary as those used for the experiment, recovered by conventional apiculture practice using a steam wax extractor. Subsequently, the wax was melted and formed into flat sheets by dipping a flat wooden form, 75 mm by 40 mm, into the molten wax. The wooden form was left to soak in water before use whereupon it was held horizontally before dipping into the liquid wax for 3 seconds. It was then removed for 15 seconds, to allow the wax to solidify, before reimmersion. Wax sheet of two thicknesses was produced by altering the number of immersions; three times produced sheets of 0.5 mm-0.6 mm thickness while six yielded sheets of 1.0 mm-1.2 mm.

#### Preparation of stimuli

The stimuli used to study comb building behaviour were formed from flat sheets of wax, 25.0 mm x 40.0 mm, placed vertically within the hive. These pieces of wax, referred to henceforth as tabs, were held in place by adhesion to the top bar of an otherwise empty frame. Plain sheets of wax were used as raw material for both the vertical backplane and tab, and as experiment specific stimuli, with the requisite pieces being prepared by cutting the sheets into strips. The strips, optionally bent as required for each experiment, were welded to the backplane using a temperature-controlled solder iron set to 205°C. A seam weld along the entire connection between a stimulus and the backplane was found to be necessary as otherwise, the bees would remove the stimulus. The exact designs used for the stimuli are detailed below.

#### Preparation of stimuli for experiment i - Initial deposition target (pit rims)

Experiment i was used to test prediction P1, that bees should initially deposit wax at the rim of a shallow dip in the wax. In this way, the walls of the finished comb will coincide with the rim of the dip. If prediction P1 is correct, the walls of cells at an early stage of construction will overlap with the rim of the seed pits significantly more often than expected by chance.

The stimuli to test the wax deposition preferences were prepared using thin sheets of wax (formed by three immersions). The resulting sheets of wax were approximately 0.5 mm thick and were cut into pieces of approximately 25 mm by 40 mm to form the tabs.

Shallow indentations were pressed into one side of a tab using a 4.0 mm-diameter domed rod. The resulting indentations were approximately 0.25 mm deep and between 3 mm and 4 mm in diameter. The obverse face showed an area where the surface texture of the wax was disturbed by compression against the support surface. The obverse face disturbance was visible and was not raised above the general surface by more than 0.1 mm. The indentations were placed *ad hoc*, 6, 7 or 8 per tab, at a distance of 10 mm to 15 mm from each other.

#### Preparation of stimuli for experiment ii - Initial deposition (pairs of pits)

Experiment ii was used to test prediction P2, that a stimulus formed from two small depressions will result in two cells conjoined at a wall aligned to the common tangent between the two pits: i.e., orthogonal to a theoretical line connecting the pit centres. If prediction P2 is correct, the alignment of a wall built between the seed pits will diverge from this common tangent less than expected by chance.

This experiment was designed to investigate the interaction between two cells that had been started close to each other, and to test prediction P2: that the stigmergy-driven reaction to adjacent shallow dips would lead to a cell wall located midway between the two stimuli.

The stimuli were prepared as described in 0, but impressions were made as pairs rather than single depressions. Pairs of shallow indentations were pressed into one side using a 4.0 mm-diameter domed rod. The indentations were from 1 mm to 3 mm apart. A total of 6-8 pairs were pressed into each sheet, at a distance of 10 mm to 15 mm from each other. The orientation of each pit pair was *ad hoc*, albeit deliberately varied. The eventual distribution of the seed orientation can be seen in Figure 1-7.

#### Preparation of stimuli for experiment iii - Initial deposition (V-form)

Experiment 3iii was used to test prediction P3: that a stimulus formed as a V-shaped barrier will result in a cell on either side of the apex conjoined at a wall that will be aligned to the bisection of the barrier. If prediction P3 is correct, the alignment of a wall built near the apex will diverge from the V shape bisection less than expected by chance. Furthermore, the wall will be positioned closer to the apex than expected by chance.

The test pieces for this experiment were fabricated using a 0.5 mm-thick backplane of ∼25 mm by 40 mm, onto the face of which 4 seed strips were added. The seed strips were 2 mm to 3 mm in height and 0.5 mm in width, prepared by cutting strips from a similar flat sheet of wax. A single strip was folded to form the V-shape and then welded onto the wax backplane. The angle of the ‘V’ was deliberately, but manually, varied, with the resulting angles to be determined by measurement from the photographic record, rather than by production control. The resulting distribution of the V angles is presented in Figure 1-10. The orientation of each V-shape was *ad hoc*, albeit deliberately varied. The eventual distribution of the seed orientation can be seen in Figure 1-8. The test pieces were bonded to the top bar of a frame using molten wax, and this frame was then inserted into a hive.

Test frames were kept in the hives for a single session, of repeated sessions, until cell construction had started, but were removed before the cells had reached a depth of 5mm. A tab carrying stimuli of this form would typically comprise one component of a test frame that would include two additional tabs presenting other stimuli.

#### Preparation of stimuli for experiment iv - Initial deposition, V-form with dual pits

Experiment ii was used to test prediction P2, that cell construction would be guided by pairs of pits and experiment iii was used to test prediction P3, that cell construction would be guided by V-shaped stimuli. This experiment, iv, was used to test whether cell construction would be guided more by one stimulus than the other by offering both stimuli but in conflicting alignments. If construction is preferentially guided by one stimulus, then wall alignment divergence from the V shape bisection will differ from its divergence from the pit common tangent by more than expected by chance.

The test pieces fabrication began using the technique described in 0 to produce V-shaped stimuli bonded to a backplane. A pair of shallow indentations were then pressed into the face of the backplane, one on either side of the apex. The alignment of the two pits was manually varied ensuring that the pit common tangent did not align with the ‘V’ bisection line.

#### Stimulus handling and construction time

Experiment-specific wax stimuli were fixed to the underside of the top bar of hive frames using molten wax. Each frame typically comprised three separate tabs, each presenting different stimuli. The frames were then placed into honeybee hives between 9 and 10 am, and were positioned at the rear limit of the brood space as this location was most likely to attract comb building activity. The test frames were inspected and photographed between 1 and 2 pm and, if necessary to achieve the requisite degree of comb construction, reinserted and removed again between 5 and 6 pm. Frames were not left in the hives overnight as the extended period of some 16 hours would likely result in excessive progress being made without the opportunity to inspect and record any interim stages of construction. All frames were removed before the cells had reached a depth of 5 mm. This process of periodic inspection allowed for progress to be monitored despite differences in the rate of comb construction. This varied between hives, but also from day to day depending on the weather and foraging opportunities.

At each inspection, the state of each frame was photographically recorded. These photographs were used to analyse the progression of comb construction, as described in section 1.3.2.

## Measurement methods

### Recording and photography

Frames were photographed before being placed in the hives and each time they were removed. This sometimes occurred only once before construction had sufficiently progressed, but often the frame would be returned to the hive and subsequently removed and re-photographed.

Photography was performed in a room adjacent to the apiary using natural daylight. The frame was mounted for each photograph on a jig that also held the camera. The jig held the camera and frame in repeatable relative positions thus the frames appear in each photograph to be of a similar size and position within the tolerances of the camera and frame mounting.

The camera (Samsung Galaxy Camera 2 EK-GC200) yielded images of 4608 by 2592 pixels. The jig held the frames 390 mm from the camera, resulting in the frame (width 333 mm) occupying an image width of approximately 3000 pixels. Image resolution was therefore approximately 9 pixels per mm.

### Photographic record analysis

Image manipulation was performed using my custom software, FormImageCompare. This tool included features for alignment of images taken before and after a treatment, magnification, marking of features, measurement of position, location and angles from those marks and other functions mentioned elsewhere in this document. FormImageCompare was written in C++ using Microsoft Visual Studio Community 2019: Version 16.7.2, Visual C++ 2019, drawing on support from the library OpenCV:Version 3.3. This tool is available at https://github.com/VinceGalloQMUL/honeycombThesisRepo.

The jig used to photograph the frames provided some consistency between photographs, but the difference from one to another was too great to allow direct comparison of wax features of the scale of cells and cell walls. This limitation was overcome by alignment during the analysis. The frames used for mounting the stimuli were marked on both sides using four crosses, one at each end of the top bar and another at each end of the bottom bar.These marks were used during the measurement stage of the analysis (0) to align the images to a degree of precision necessary for the comparison of features such as cell walls and wall junctions. Alignment of the image pairs was achieved using FormImageCompare to align the images.Alignment markers were placed by mouse-click when the pointer was the centre of a mark. Four such alignment points featured located for each image using crosses pre-emptively marked on the test frames. The software could then align the locations on the second image with those on the first (using the functions calcPerspectiveLambda(..) followed by perspectiveTransform(..) contained in the library OpenCV) eliminating displacement of the image or perspective change between first and second photograph.

Once the two images had been aligned it was possible adjust their relative contrast to view one image, or the other, or a blend of both. In this way, the position of a comb feature could be compared with either the stimulus feature that had initially been offered to the bees, or with the condition of the feature at some earlier stage of construction.

The software included features whereby the user could place additional feature markers to identify locations allowing specific measurements to be made, each of relevance to an individual cell feature. The detail of each experiment specific measurement is described in the relevant section below.

### Experiment-specific measurements

#### Experiment i - Initial deposition target (pit rims)

To determine the degree of overlap between the rims of the seed pits and the eventual comb built on each wax panel, I analysed photographs of the tabs at the start and end of the experiment; recording the positions of pit rims at the start and cell walls at the end.

The measurements obtained from the experiments provided the sample distribution. A control sample was generated by the randomised placement of pseudo-depressions, for which the overlap with whatever cell walls lay beneath it was measured.

Measurements of the location of pits and the associated cell wall used the coordinates of points marked on paired images, taken of each face of the subject comb before and after it had been built. The photographs recorded each side of the test pieces prior to their placement in the hives, and also at each stage of inspection. The frame comparison tool FormImageCompare includes a feature to measure ‘pitRim’. This feature allows the user, when viewing the first image recorded before any comb had been built, to mark the centre of a depression made into the tab, whereupon the software draws a blue circle scaled to the equivalent of 4.0 mm diameter.

Using the second image, which displayed the state of the item after some comb had been built, the user could mark the locations where the guide circle (representing the rim of the pit) overlapped the walls that had been built. Each overlap was marked by clicking the start intersection between the circle and the wall, followed by a subsequent click to set the location where the overlap ceased. The overlapping chord was drawn as a white line by the software together with a small circle, which marked the starting end. The start and end of a chord were assumed to be placed in a clockwise direction, which allows the software to unambiguously measure the angle subtended by the chord. The software calculated the total angular overlap for each pit by summing all chord marks created by the user.

The layout of a pit and surrounding cell that was marked by the operator on each occasion will be of the form shown in Figure 1-1.

**Figure 1-1.**
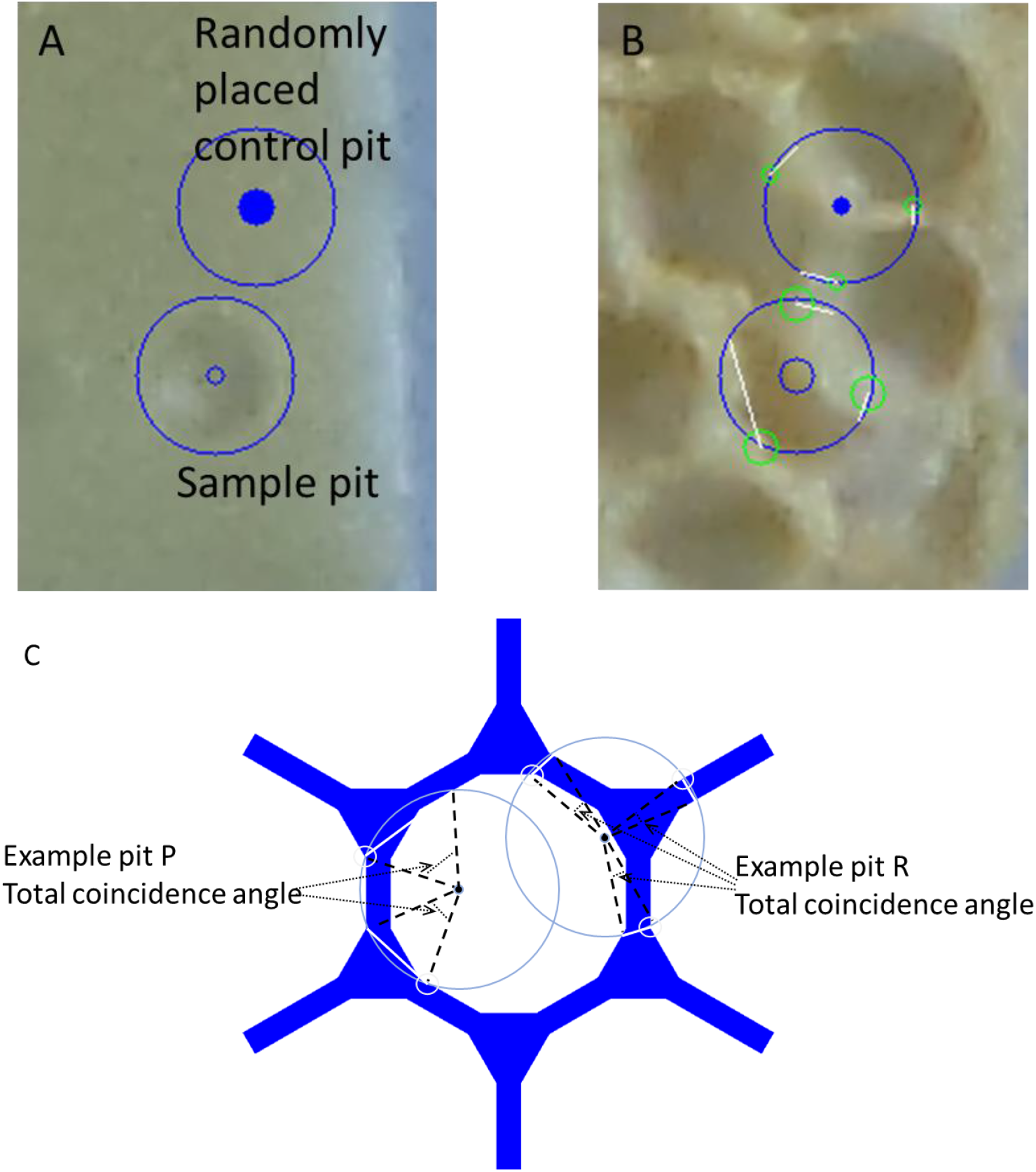
Pit rim to cell wall coincidence measurement. A: both a sample pit, visible as a depression in the wax, around which the software has drawn circle to act as a measurement gauge, and a software-generated randomly placed control pit. B: a view after the operator had marked the extent of overlap between the gauge circles and the cell walls. C: a diagrammatic representation of the measurements to be taken from the gauge and overlap marks. The metric, calculated and exported by the software, is the total angular overlap for each pit; larger in the diagram for example pit P than for example pit R.

Samples used as a control population were randomly generated. For each pit marked by the operator, the software drew an additional circle as a pseudo-pit placed at random within 10.0 mm of the original. I then marked the start and end of chord segments where the pseudo-pit rim overlapped cell walls. The software calculated the total angular overlap for each pseudo pit in the same manner as for a manually located pit.

#### Experiment ii - Initial deposition (pairs of pits)

Measurements of the location of pits and the associated cell wall used the coordinates of points marked on paired images taken of each face of the subject comb, recorded both before and after the comb had been built, as described in 1.3.3.1. The frame comparison tool FormImageCompare includes a feature to measure ‘pitPair’. This feature allows the user, when viewing the first image recorded before any comb had been built, to mark the centres of a pair of depressions whereupon the software draws a blue circle scaled to the equivalent of 4.0 mm diameter for each pit. The location of the two centres allows the software to calculate the orientation of the line between the centres, and hence that of the orthogonal pit common tangent.

Using the second image, recorded post-construction, the user could mark the line of a wall found between the pits, or of the nearest wall if none were found between the pit centres.

The layout of a pit pair and cell wall that has been marked by the operator is shown in Figure 1-2. For each pit pair and cell wall the results comprised the orientation of the wall and that of the pit common tangent, and the divergence of these from one from another.

**Figure 1-2.**
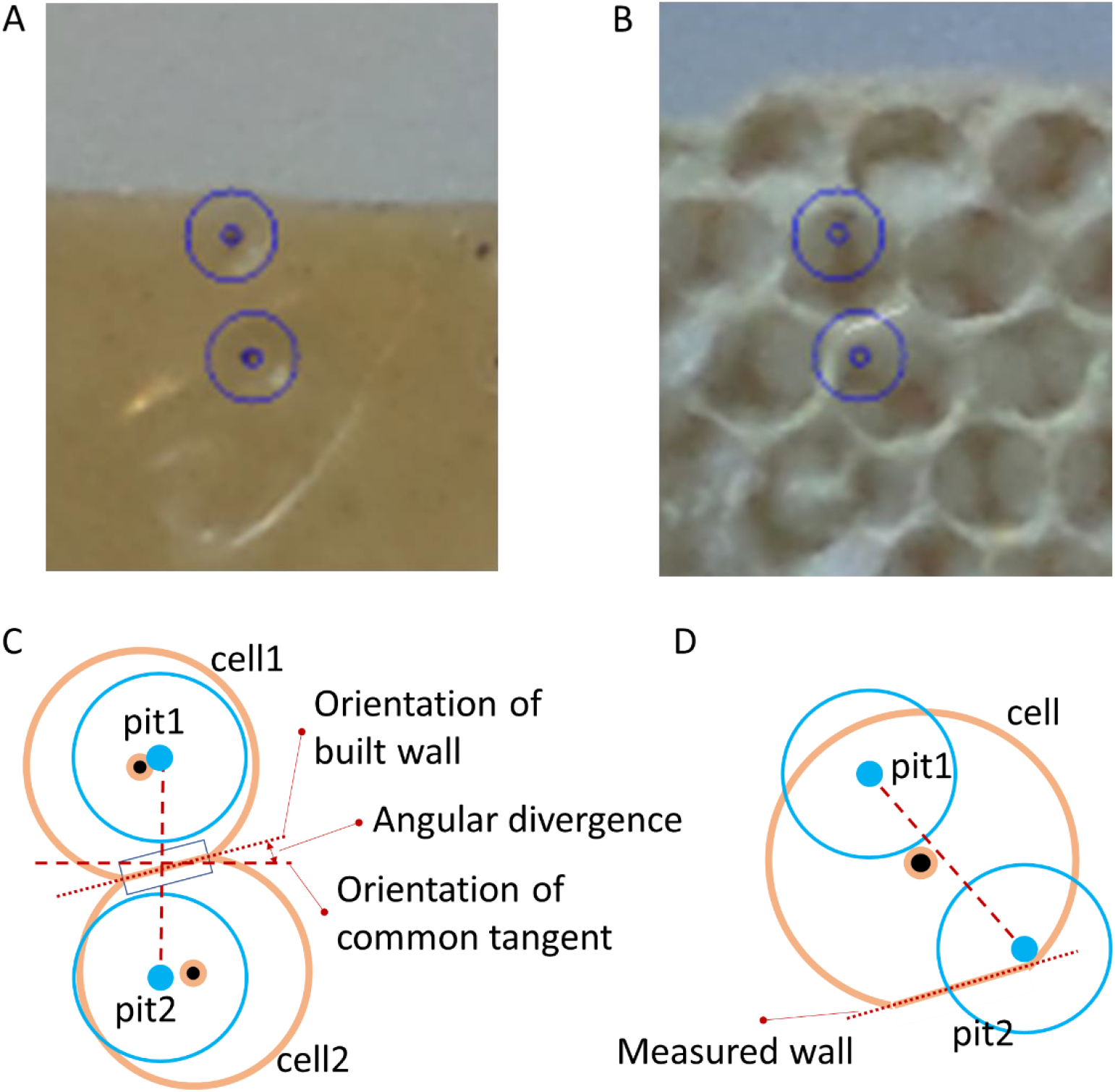
Measurement of cell wall located between pit-pair stimulus. A: the marks placed on the photograph recording the initial state of the stimulus, highlighting the location of the pits. B: the photographic record of the same tba after construction had begun, showing the location of the original pits, and the line used to mark the cell wall between the pit centres. C: diagrammatic representation of the stimulus, the resulting comb, the marks placed by the operator and the resulting angle computed by the software. D: an example of an exception where the nearest wall was not between the pit centres.

Samples used as a control population were randomly generated. The software drew an additional pair of pit marks at a random location, orientation and separation (between 4.0 and 5.0 mm). The operator then marked a wall found between the two random marks in the same fashion as previously done for real stimuli.

As described in 1.3.3.1., as before, after some comb was built on a tab, in many cases the cells formed regular structures that may have reduced the independence of separate pit/cell wall measurements obtained from a single tab. Therefore, the mean of the measurements was used to form a single measurement for each tab.

The distribution of divergences between built walls and their associated common tangents was normally distributed around 0°. However, the analysis was performed using absolute divergence which has a single-tailed distribution (Figure 1-6). Thus, comparison of the experimental and control samples was done using a Wilcoxon ranked comparison test.

#### Experiment iii - Initial deposition (V-form)

The alignment between a V-shaped seed and the associated cell wall was measured using the coordinates of points marked on paired images of each face of the subject comb, recorded before and after construction. The frame comparison tool FormImageCompare includes a feature to measure ‘bend’. This feature allows the user, when viewing the first image recorded before any comb had been built, to mark the ends of both strips that form the ‘V’ and the apex between. Together, these locations allow the software to calculate the orientation of the bisection.

Using the second image, recorded post-construction, the user could mark the line of an inter-cell wall that was closest to the apex by placing two marks at the ends of the nearest wall at corner 1 (closest to the apex) and corner 2, further in the direction of the apex, (Figure 1-3 C). Together, these corner locations provided the orientation of the wall and therefore allowed the calculation of the angular divergence between the wall and the V-shape bisection. The 2^nd^ closest wall also needed identifying and marking. Starting from corner 1, following the cell wall and passing through (or close to) the apex, to the next two cell corners (corners 3 and 4, Figure 1-3 C). The 2^nd^ closest wall is that between corners 3 and 4.

**Figure 1-3.**
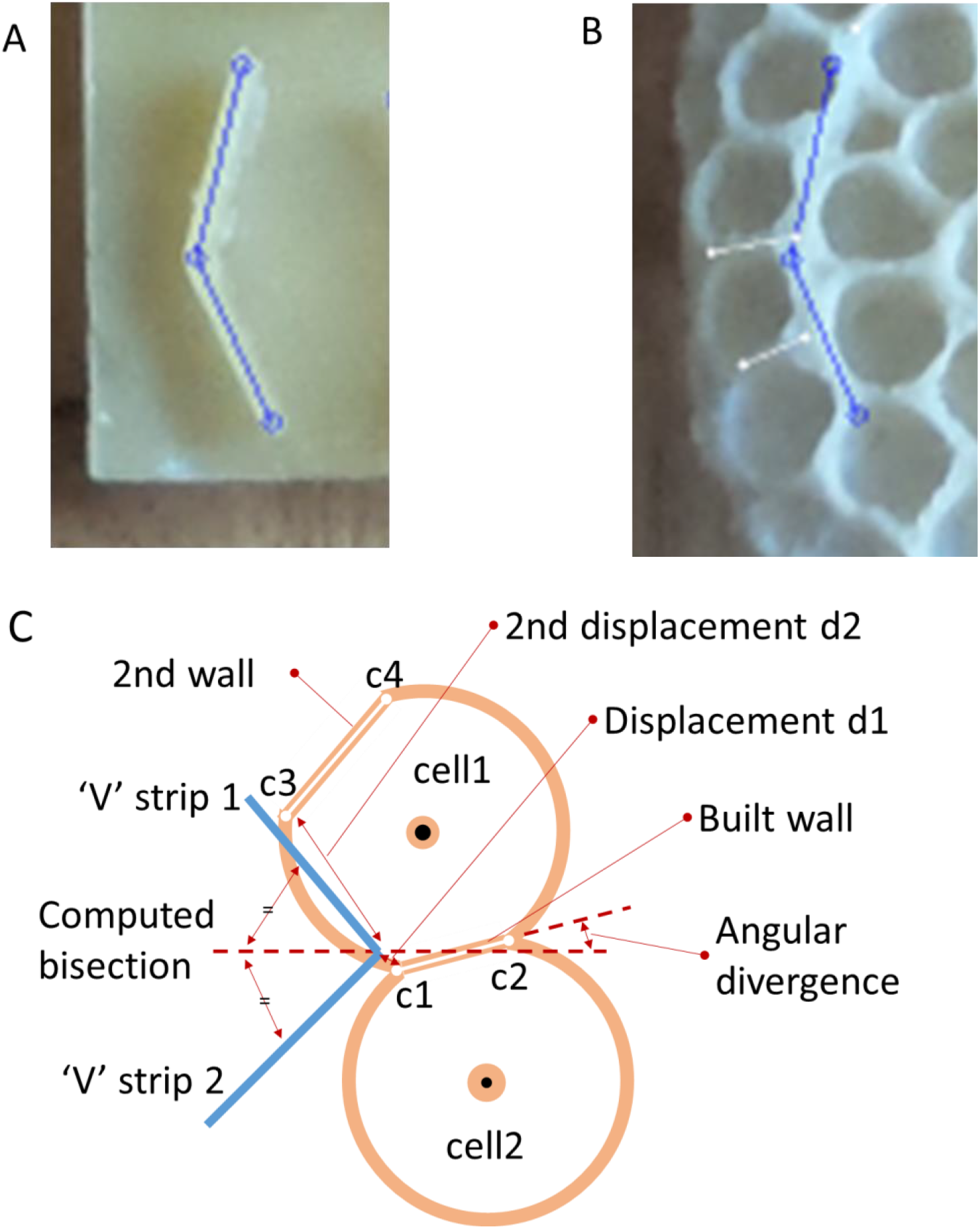
Measurement of cell wall located at the apex of a V-shaped stimulus. A: the two ends and the apex of the V-shaped stimulus marked using the earlier of the two images. B: switching to the aligned, later stage, image the two closest cell walls are shown, marked in white. C: a diagrammatic representation of the V, two cells built close by, the two closest walls. The diagram also shows the measurements, computed and exported by the software: the angle difference between the closest wall and the V bisection, and both wall (corner) displacements.

Samples used as a control population were randomly generated. The software drew an additional V mark at a random location, orientation and splay (between 90° and 152.2°). I then marked the walls associated with the random mark in the same fashion described for real stimuli.

#### Experiment iv - Initial deposition (V-form with dual pits)

For this experiment, stimuli were configured to have both a V-shaped seed and two pit depressions. Measurement of the outcome from these used the same techniques as those described for experiment 3ii and experiment 3ii. The frame comparison tool FormImageCompare includes a feature to measure ‘bend’. This feature allows the user, when viewing the first image recorded before any comb had been built, to mark the ends of both strips that form the ‘V’ and the apex between. This feature also allows the user to mark the centres of a pair of depressions. The software then confirms the placement of the marks by drawing circles for each mark, with lines joining the ‘V’ locations.

The second image, recorded post-construction, was used to mark the line of an inter-cell wall closest to the apex. Two marks placed at the ends of the nearest wall provide the orientation of the wall and, therefore, allow the calculation of the angular divergence between the wall and the V-shape bisection, as well as the divergence between the wall and the pit common tangent.

The ‘V’ stimulus marks allowed the software to calculate the orientation of the bisection line and, from the locations of both pit centres, to calculate the orientation of the pit common tangent. Comparing these values with the orientation of the built wall yielded the divergence of the wall orientation from each of the seed stimuli.

The layout of a V seed, the pit pair and the cell wall that was marked by the operator on each occasion is shown in Figure 1-4.

**Figure 1-4.**
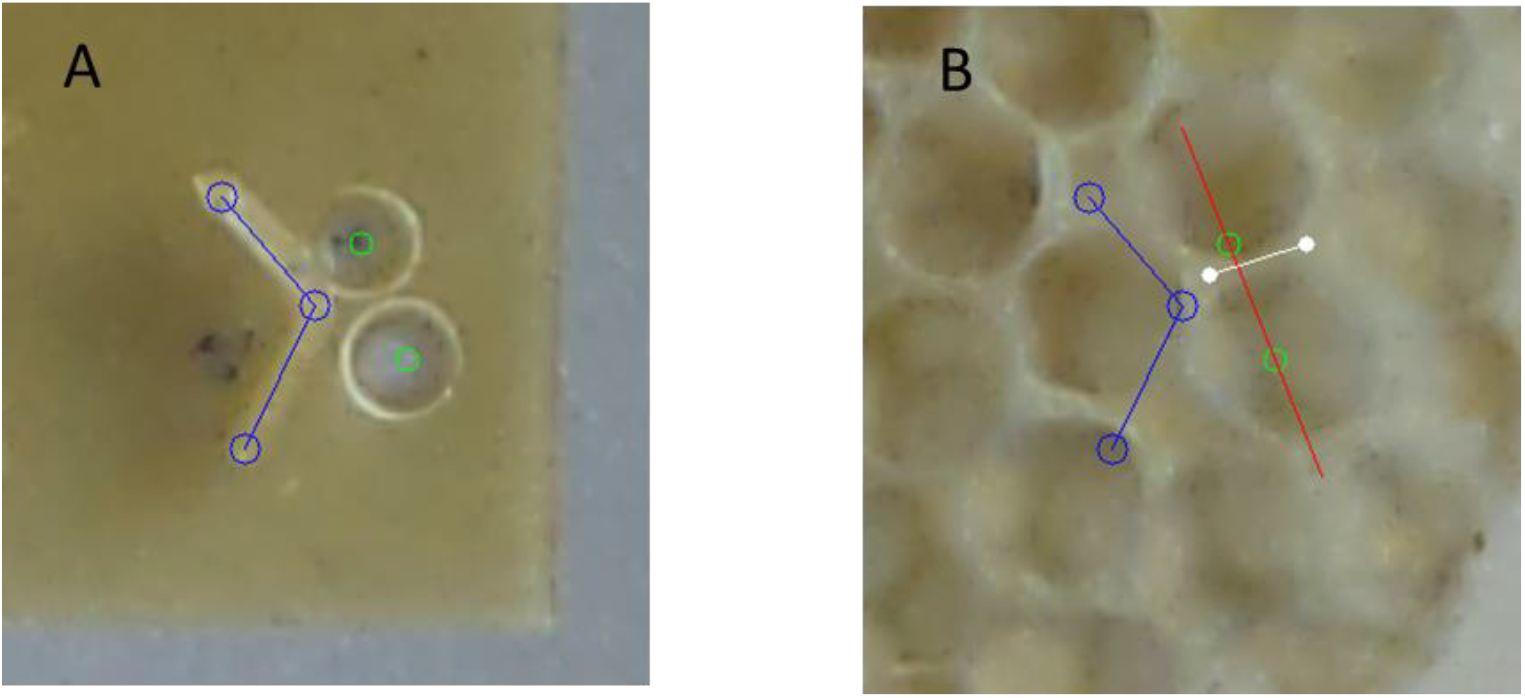
Measurement of cell wall located at the apex of a combined stimulus formed of a V-strip and pit-pair. A: marking the early-stage record at the two ends and the apex of the V and the centre of both pits. B: switching to the aligned, later stage, image the cell walls closest to the V apex is marked in white. The later image includes the computer drawn marks locating the initial compound, stimulus.

## Analysis

Data obtained from the images, using FormImageCompare (0), was subsequently process using custom scripts written in R and run within RStudio version 1.3.1093 incorporating R version 3.6.3.

Each experiment offered to the bees flat wax backplanes, tabs, upon which stimuli were mounted or imprinted. Each tab carried multiple stimuli resulting in multiple measurements from each tab. While the cell pattern built onto some of the stimuli were limited in scope, not reaching to adjacent stimuli, some tabs had comb built to cover much of the surface.

The regular layout of cells within such larger patched of comb may introduce a degree of interdependence of results obtained for separate pit/cell wall measurements obtained from a single tab. The mean of measurements obtained from each tab formed a single measurement per tab.

### Experiment i - Initial deposition target (pit rims)

The measurement for each sample was the angular overlap between the rim of the pit stimulus and the wall of subsequent cells. This measurement was made for both physical pits offered as stimuli, and for randomly placed pseudo-pits. Comparison between experiment and random populations was made using a T-test, using the R function t.test(), as the data appear to be normally distributed (Figure 1-5).

**Figure 1-5.**
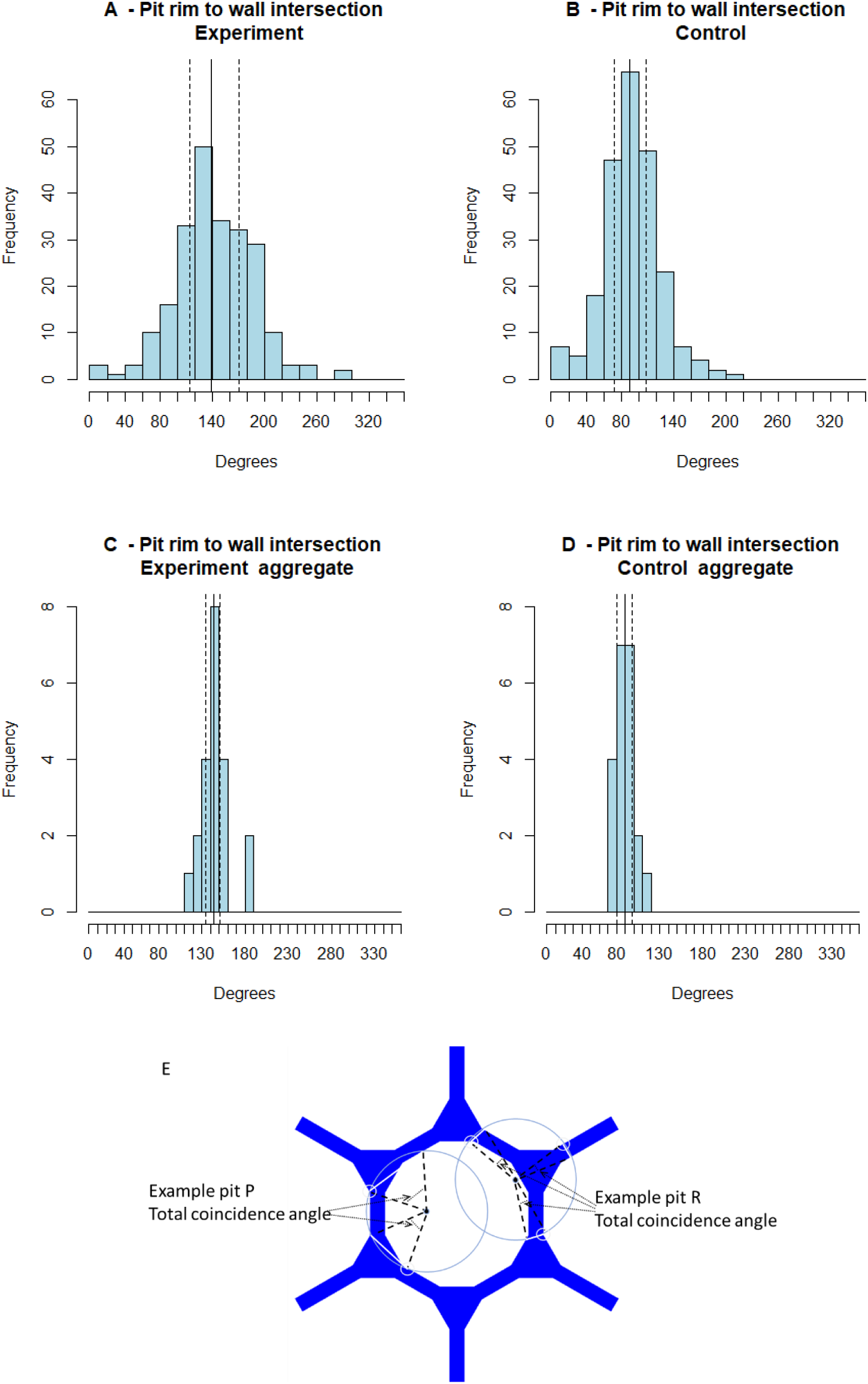
Normalised distribution of the angular overlap between pit rim and cell walls. All charts include a solid line showing the median value and dashed lines indicating the 1^st^ and 3^rd^ quartiles. A: shows the distribution of all experimental samples. B: distribution of values the complete set of control samples. C: distribution of experimental overlaps obtained by averaging over each tab. D: distribution of overlaps for the control set after averaging for each tab. E: diagram showing, theoretically, how the walls of a cell may overlap each of two pits, the total angle of which forms the metric for this experiment.

### Experiment ii - Initial deposition (pairs of pits)

The measurement made of each wall between a pit-pair was the angular difference between that wall and the theoretical tangent common to the two pits. This measurement was made for both physical pits offered as stimuli, and for randomly placed pseudo-pits. Comparison between experiment and random populations was made using the R function wilcox.test(), a Wilcoxon ranked test, as the data are unbalanced measuring divergence (>0) rather than angular difference (+ve and -ve) and appear not to be normally distributed (Figure 1-6).

**Figure 1-6.**
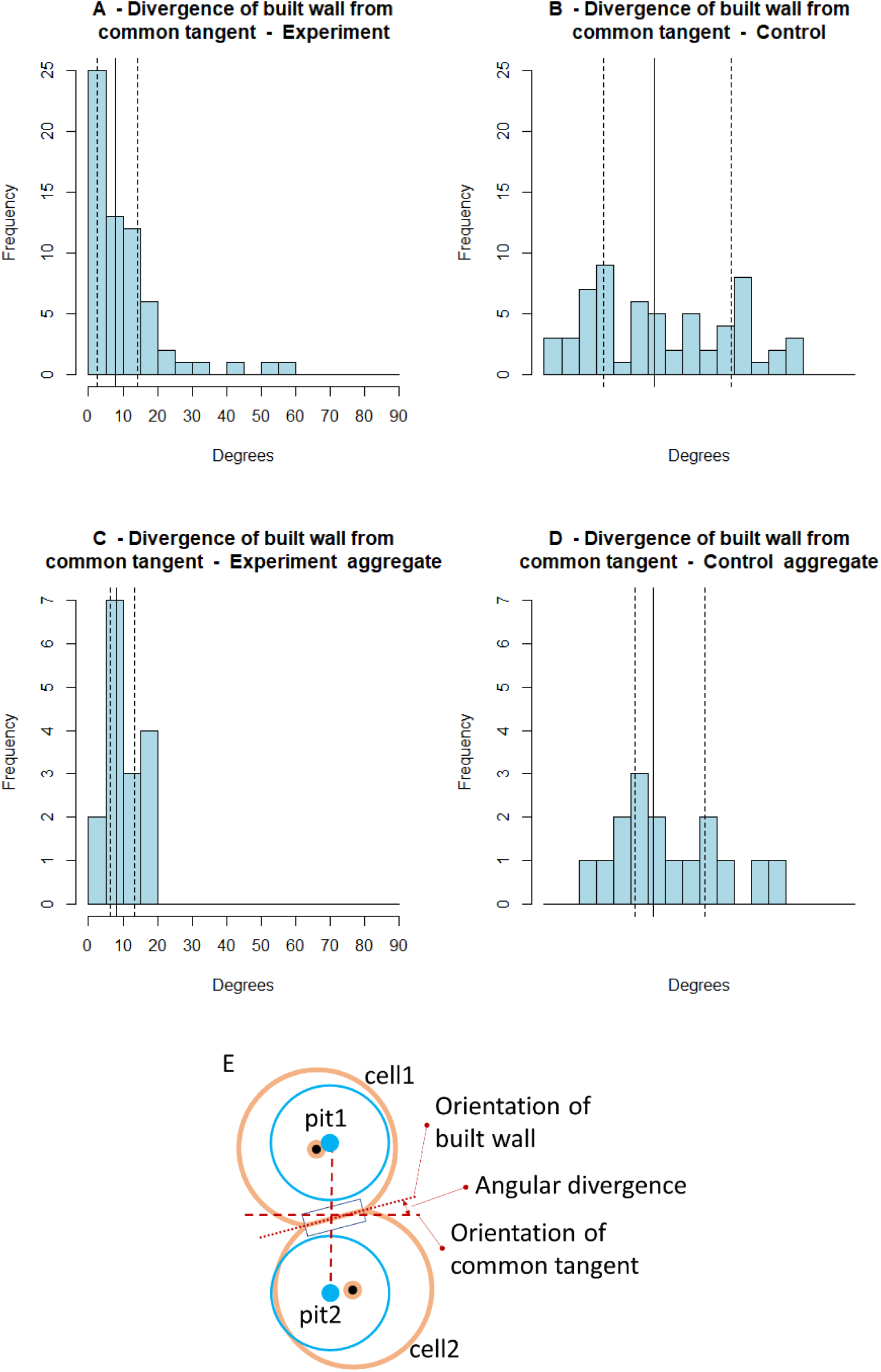
Distributions of divergence between the orientation of the constructed wall and the common tangent stimulus. All charts include a solid line showing the median value and dashed lines indicating the 1^st^ and 3^rd^ quartiles. Chart A shows the distribution of all experimental samples while B shows that for the complete set of control samples. The values used for the statistical analysis were those obtained by averaging over each tab. The distribution of the aggregated divergences is shown in C for the experimental set and the control set in D. E: diagram showing, theoretically, how the relationship between pits, cells and the relevant built wall and its divergence from the common tangent which forms the metric for this experiment.

### Experiment iii - Initial deposition (V-form)

Two measurements made of cell walls close to the apex of V-form stimulus. The first measurement was the angular difference between that wall and the theoretical line bisecting the V-form. The second measurement pair was the distance, d1, from the V-form apex to the closest cell corner and the distance, d2, from the apex to the next nearest corner (Figure 1-3 C). These distances were combined to form a metric of proximity, P; the ratio calculated as P = d1 / (d1 + d2). Comparison between experiment and random populations, for both metrics, was made using the R function wilcox.test(), a Wilcoxon ranked test, as the data appear not to be normally distributed (Figure 1-9 & Figure 1-11).

## Results

### Experiment i - Cell wall position was influenced by pit placement

Measurements were taken from 14 frames, each of which carried 3 wax sheets into which the pit depression stimuli had been pressed. A total of 233 pits and the subsequent beginnings of comb cells were identified and measured on 21 tabs. A further 233 pseudo pits were created from which the random control values were obtained. Data are presented as the mean ± standard deviation throughout.

When some comb was built, cell walls overlapped with the rims of the seed pits by 144.9°±17.6°, which was significantly greater than the overlap with randomly placed pseudo pits (89.4°±11.2°; t_20_ = 11.9, P <0.00001; Figure 1-5 C&D)This demonstrated that pit placement influenced the positions of cell walls, as predicted by P3.1 – that wax deposition will begin at the edges of a stimulus comprising a shallow depression, leading to the eventual location of cell walls - when one assumes that upon encountering a sub-cell sized concave shape, a builder’s reaction will be to extend the depression by excavation of wax from the centre.

### Experiment ii – Cell wall placement was influenced by pit-pair placement

Measurements were taken from 16 tabs carrying pairs of pits. A total of 66 such pairs and the subsequent beginnings of comb cells were identified and measured. A further 66 pseudo pits were created from which the random control values were obtained. For three of these from the experimental set and five from the control set, the built wall extended beyond the centres of the seed pits and so these were excluded. Analysis was applied to the remaining 63 and 61, respectively. Data are presented as the mean ± standard deviation throughout.

When some comb was built, cell walls diverged from the pit common tangent by 9.8°±5.1°, which was significantly less than the divergence from the common tangent of randomly placed pseudo pits (35.9°±15.5°; W = 8, P < 0.00001; Figure 1-6 C&D). This demonstrates that pit placement influenced the positions of cell walls, as predicted by P3.2, that a stimulus formed from two small depressions will result in a wall aligned to the common tangent between the two pits.

For the experimental set, the angle at which the common tangent lay (clockwise angle from horizontal) compared with that of the orientation of the built cell wall is shown in Figure 1-7. This illustration shows the values for all occurrences of the pit pair measurements.

**Figure 1-7.**
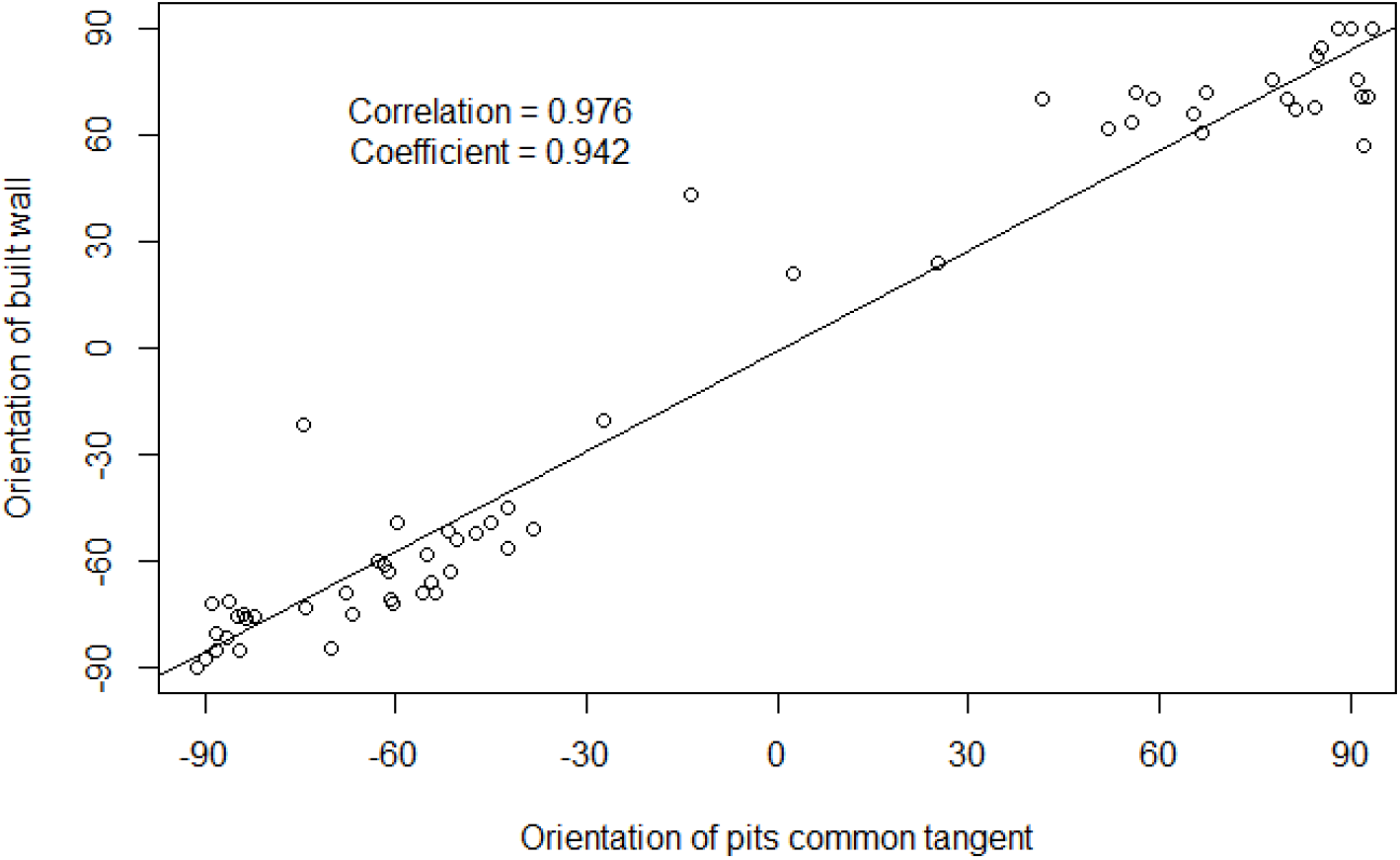
Relationship between the orientation of the built wall and that of the pit common tangent. The measurement of orientation is based on zero degrees being horizontal. The value for the common tangent is plotted on the X-axis, with that for the built wall plotted on the Y-axis. The results show a strong correlation between the two attributes, as well as a relational coefficient close to unity.

### Experiment iii - Cell wall position was influenced by V-strip placement

Measurements were taken from 16 tabs carrying ‘V’ stimuli. A total of 79 such stimuli and the subsequent beginnings of comb cells were identified and measured. A further 76 pseudo-V stimuli were also created, from which the random control values were obtained.

When some comb was built, cell walls diverged from the ‘V’ bisection by 6.4°±5.7°, which was significantly less than the divergence from the bisection of randomly placed ‘V’s (18.9°±14.8°; W = 8, P < 0.00001; Figure 1-9 C&D). This demonstrates that ‘V’ strip placement influenced the positions of cell walls, as predicted by P3.3, that each arm of the V-shape will promote formation of a cells resulting in a cojoined wall at the apex.

The correspondence between the orientation of the ‘V’ bisection and that of the built wall is shown in Figure 1-8.

**Figure 1-8.**
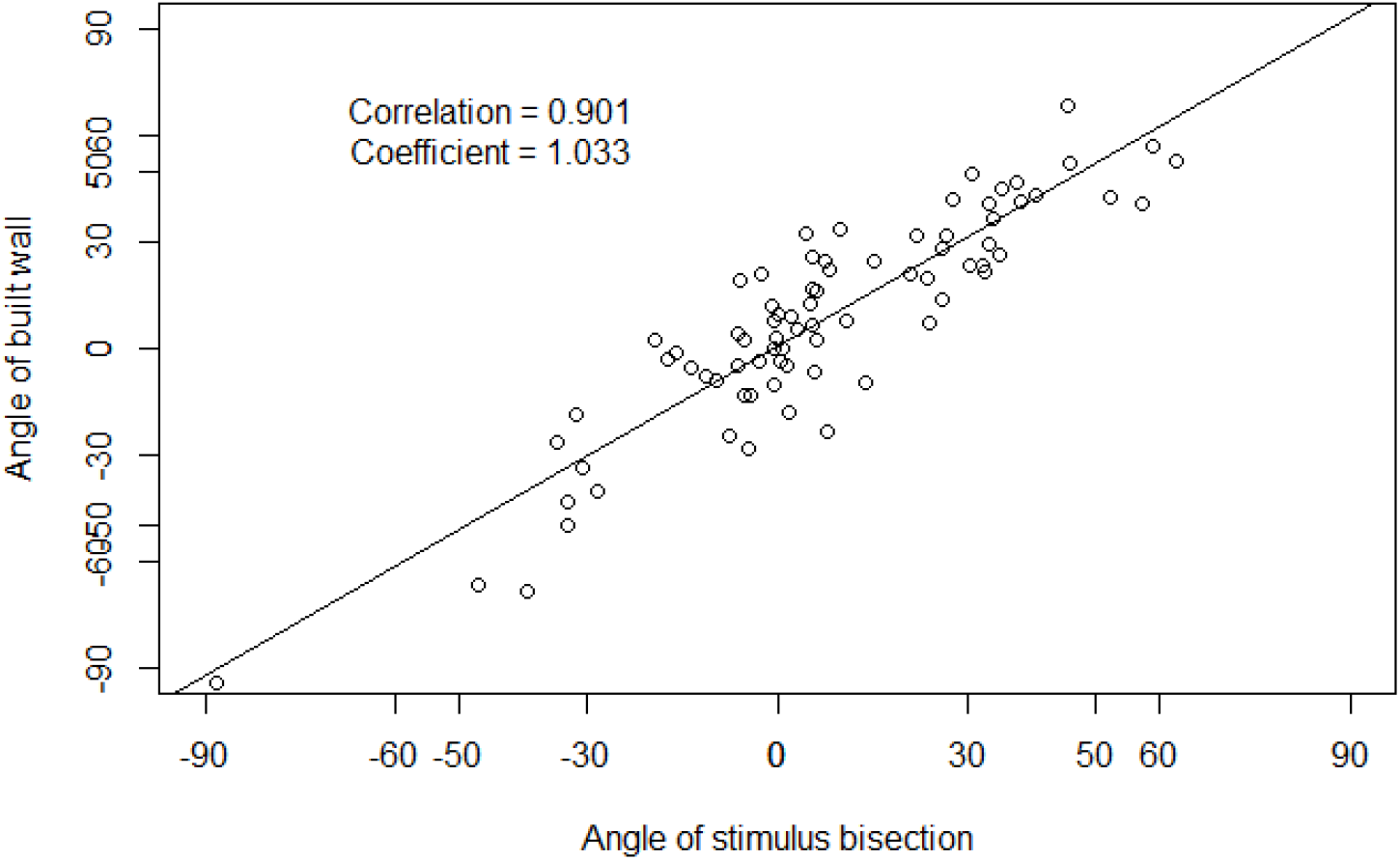
Relationship between the orientation of the built wall and that of the ‘V’ bisection. The measurement of orientation is based on zero degrees being horizontal. The value for the ‘V’ bisection is plotted on the X-axis, with that for the built wall plotted on the Y-axis. The results show a strong correlation between the two attributes, as well as a relational coefficient close to unity.

#### Proximity of wall to the apex

Measurements were taken from 16 tabs carrying ‘V’ stimuli. A total of 79 such stimuli and the subsequent beginnings of comb cells were identified and measured. A further 76 pseudo-’V’s were also created, from which the random control values were obtained.

When some comb was built, the distance between the ‘V’ apex and the nearest wall, as a fraction of the distance from that wall to the next nearest was 0.18±0.08, which was significantly less than that measurement for the apex of randomly placed ‘V’s (0.37±0.06; W = 7, P < 0.00001; Figure 1-11 C&D). This demonstrated that ‘V’ strip placement influenced the positions of cell walls, as predicted by P3.3, that each arm of the V-shape will promote formation of a cells resulting in a cojoined wall at the apex.

### Experiment iv - Cell wall position was preferentially influenced by pit-pair placement

Measurements were taken from 81 V-shaped stimuli adorned with pits.

When comb was built on the combined ‘V’ and pit stimuli, 55 built walls, from a sample of 81, were closer to the alignment of the pit common tangent than to the ‘V’ bisection (Figure 1-12). The cell walls diverged from the pit common tangent by 8.9°±8.7°, which was significantly less than the divergence from the ‘V’ bisection (14.0°±9.2°, P = 0.00005; Figure 1-13 A&B). This demonstrated that pit placement influenced the positions of cell walls more than the ‘V’ strip.

Comparing the divergence from the pit common tangent with or without the additional ‘V’ stimulus, this experiment (8.9°±8.7°, Figure 1-13 A), was not significantly different from experiment 3 (10.3°±11.7°, Figure 1-9 A, also Figure 1-13 C ;P = 0.72). This demonstrated that in the presence of pits, the ‘V’ strip placement had no more influence over the positions of cell walls than may be expected by chance.

**Figure 1-9.**
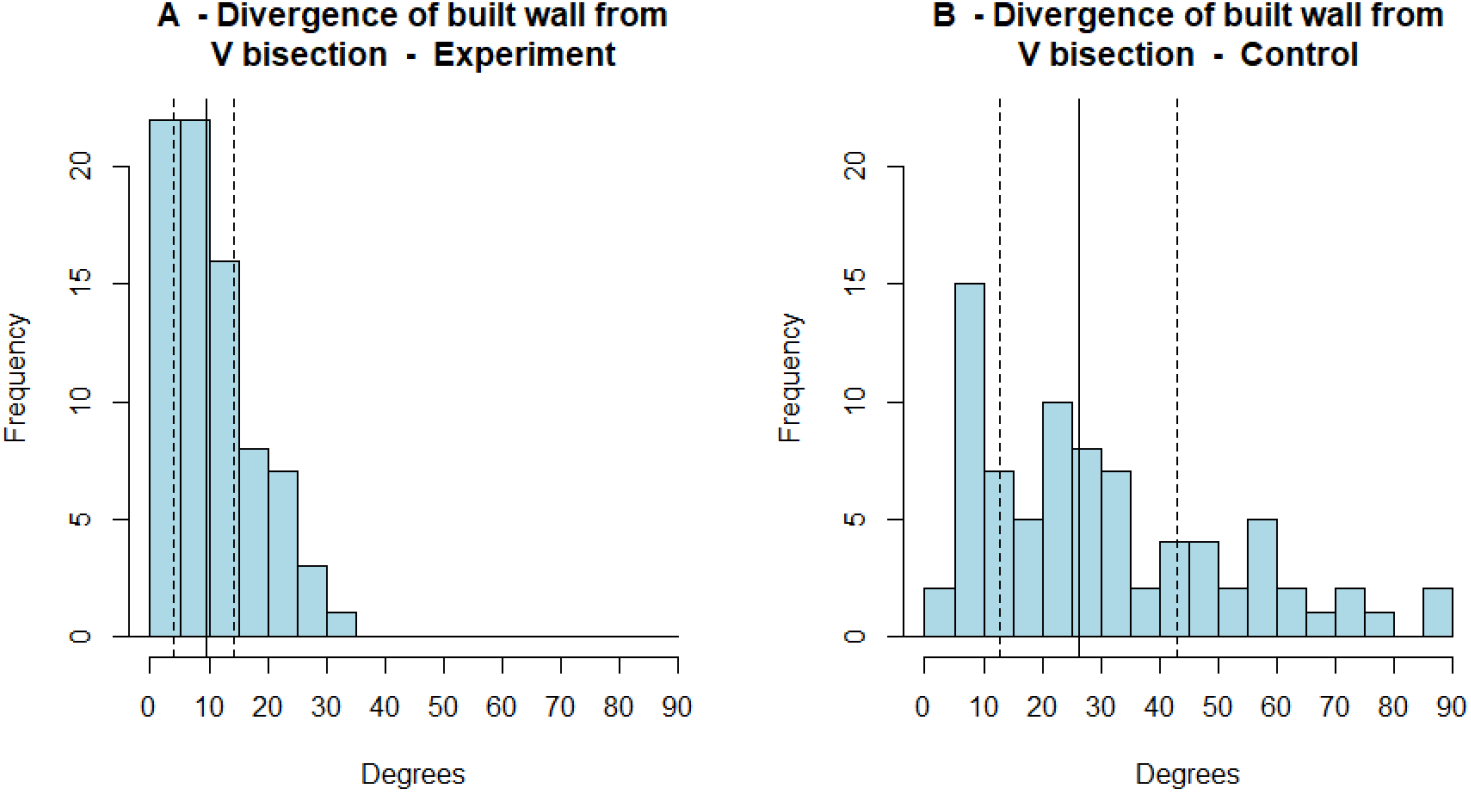
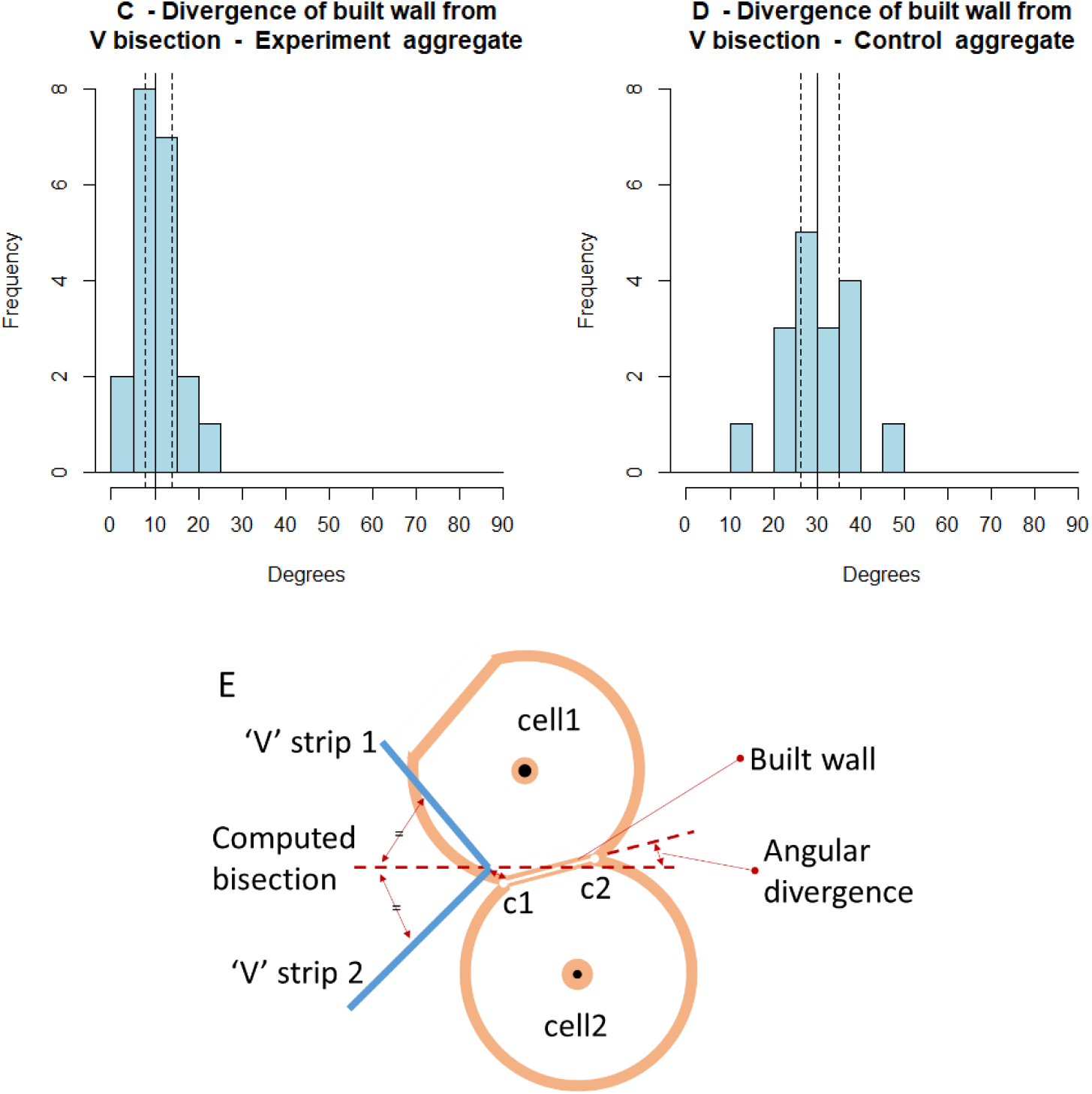
Distributions of divergence between the orientation of the constructed wall and the ‘V’ stimulus bisection. Chart A shows the distribution of all experimental samples while B shows that for the complete set of control samples. The values used for the statistical analysis were those obtained by averaging over each tab. The distribution of the aggregated divergences is shown in C for the experimental set and the control set in D. E: diagram showing, theoretically, the relationship between ‘V’, cells and the relevant built wall and its divergence from the ‘V’ which forms the metric for this experiment.

**Figure 1-10 -.**
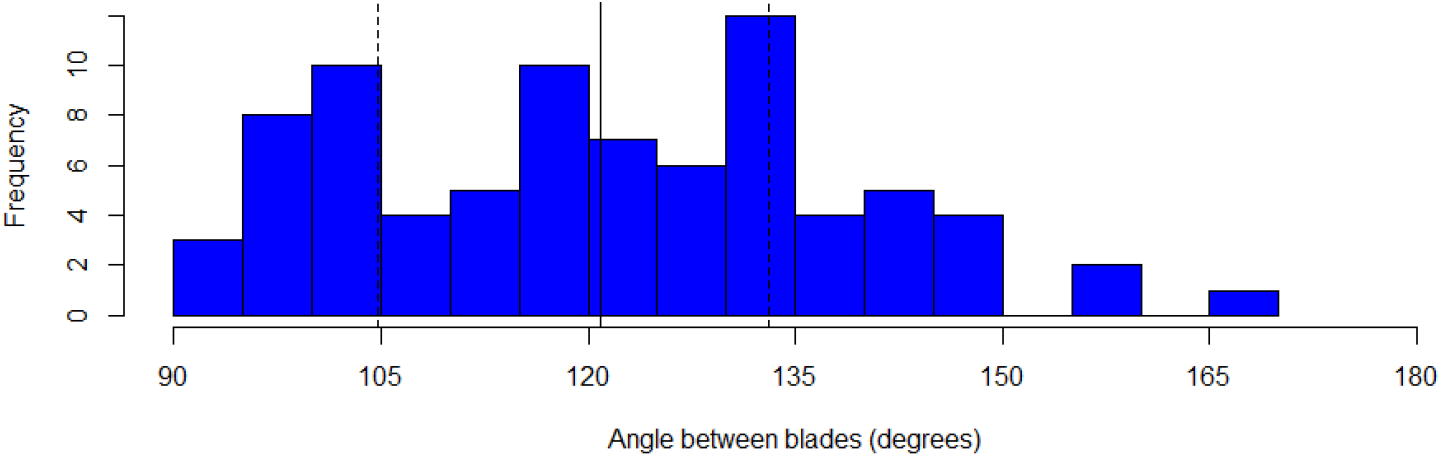
Distribution of apex angular splay for the V-shaped stimuli. The stimuli were fashioned by hand to have a distinct bend, but not less than 90°. This histogram of the ‘V’ angles, acquired as part of the measurement process, shows their distribution to be mostly from 90° to 150°.

**Figure 1-11.**
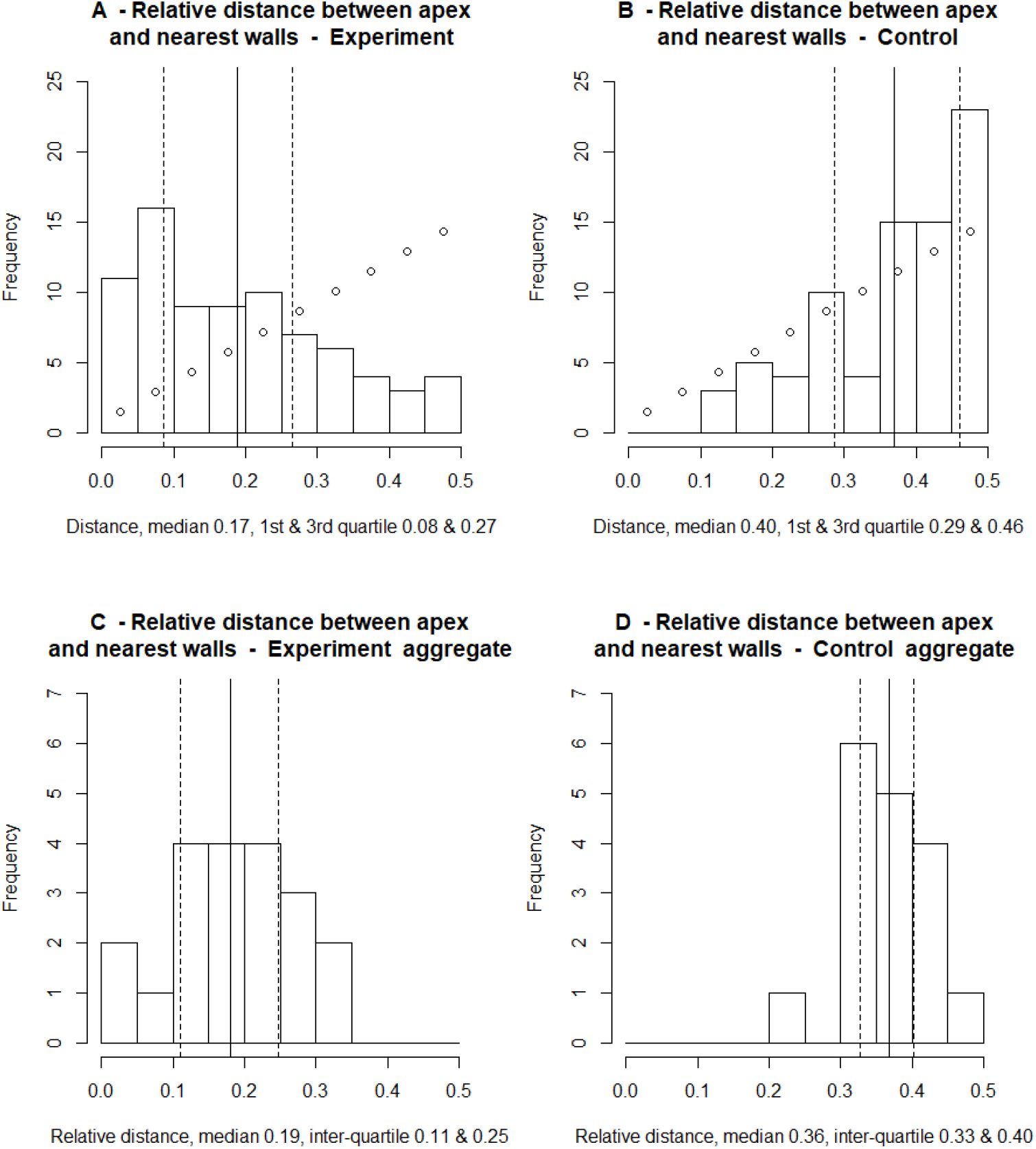
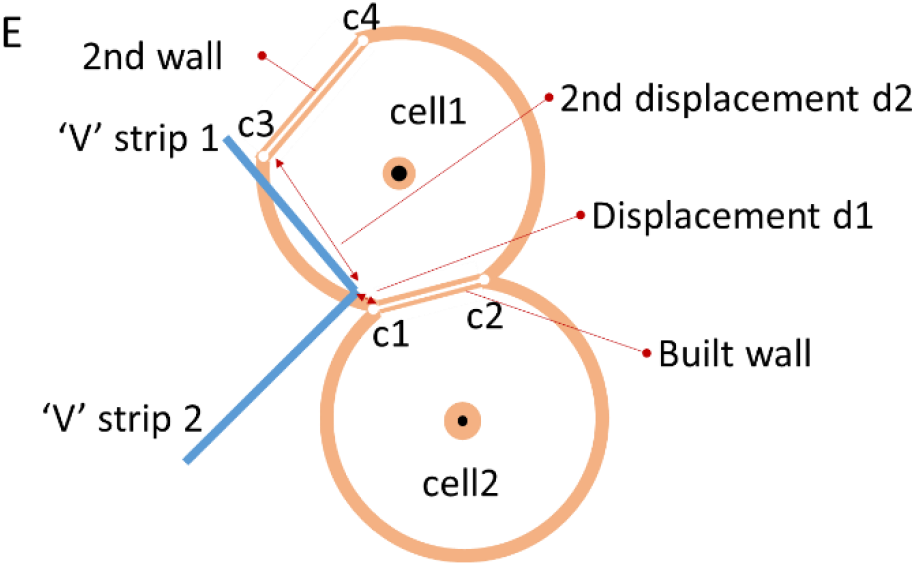
Distributions of distances between the apex of the ‘V’ stimulus and the nearest cell corner, expressed as a fraction of the distance to the nearest and the next nearest corner. Chart A shows the distribution of all experimental samples while B shows that for the complete set of control samples. The values used for the statistical analysis were those obtained by averaging over each tab. The distribution of the aggregated divergences is shown in C for the experimental set and the control set in D. E: diagram showing, theoretically, the relationship between ‘V’, cells and the relevant built wall proximity values which forms the metric for this experiment.

**Figure 1-12.**
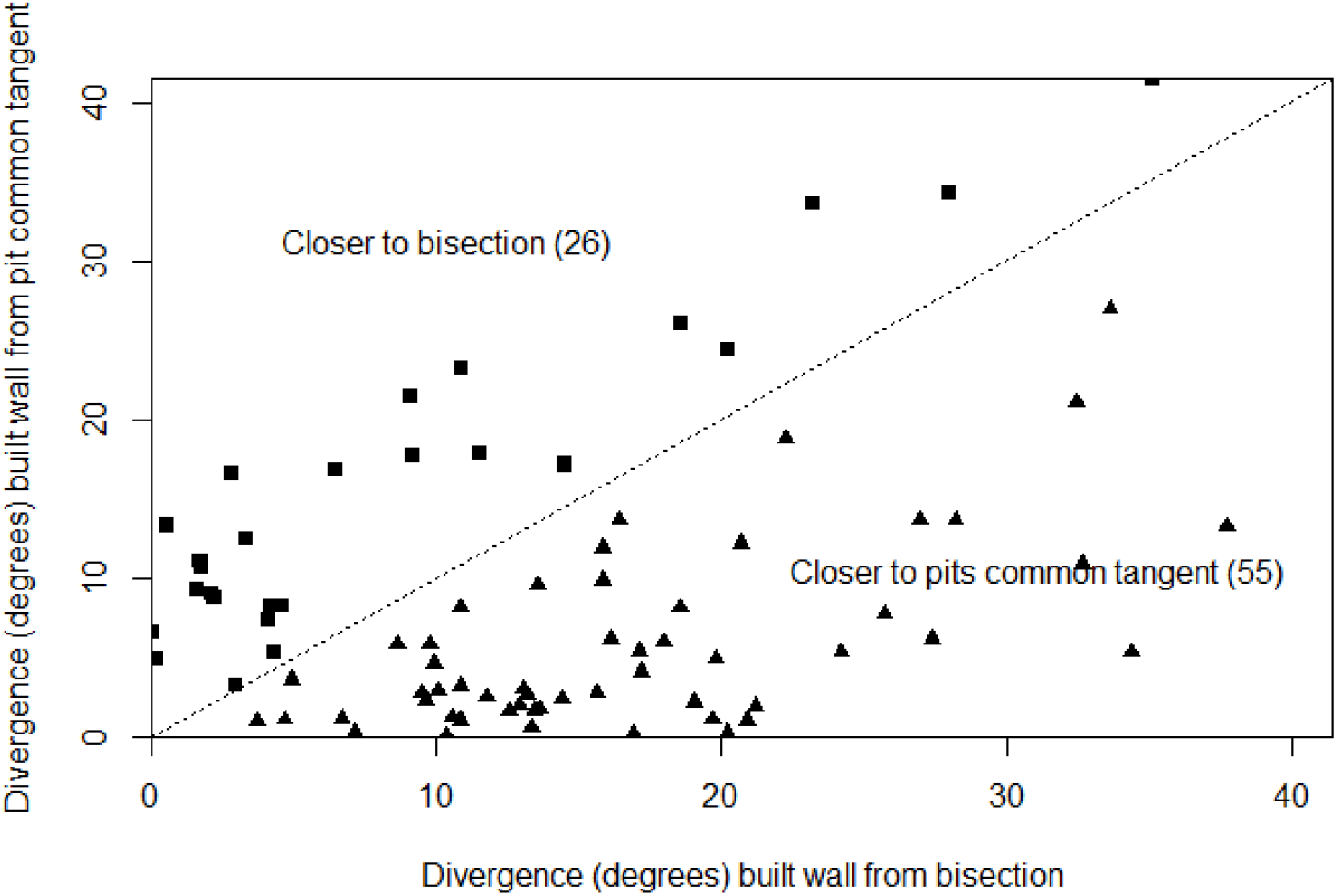
Divergence of the built wall from each of the two potential guides; V bisection and pit common tangent measured for hybrid stimuli. The division line separates the graph area into regions closer to one influence than the other. Of the population of 81, 55 walls (shown as triangles) were aligned more closely to the pit common tangent compared with 26 (shown as squares) aligned closer to the V bisection line. Alignment of the pits had a greater influence that the V-shape had on the orientation of more than 2/3^rd^ of the measured walls.

**Figure 1-13.**
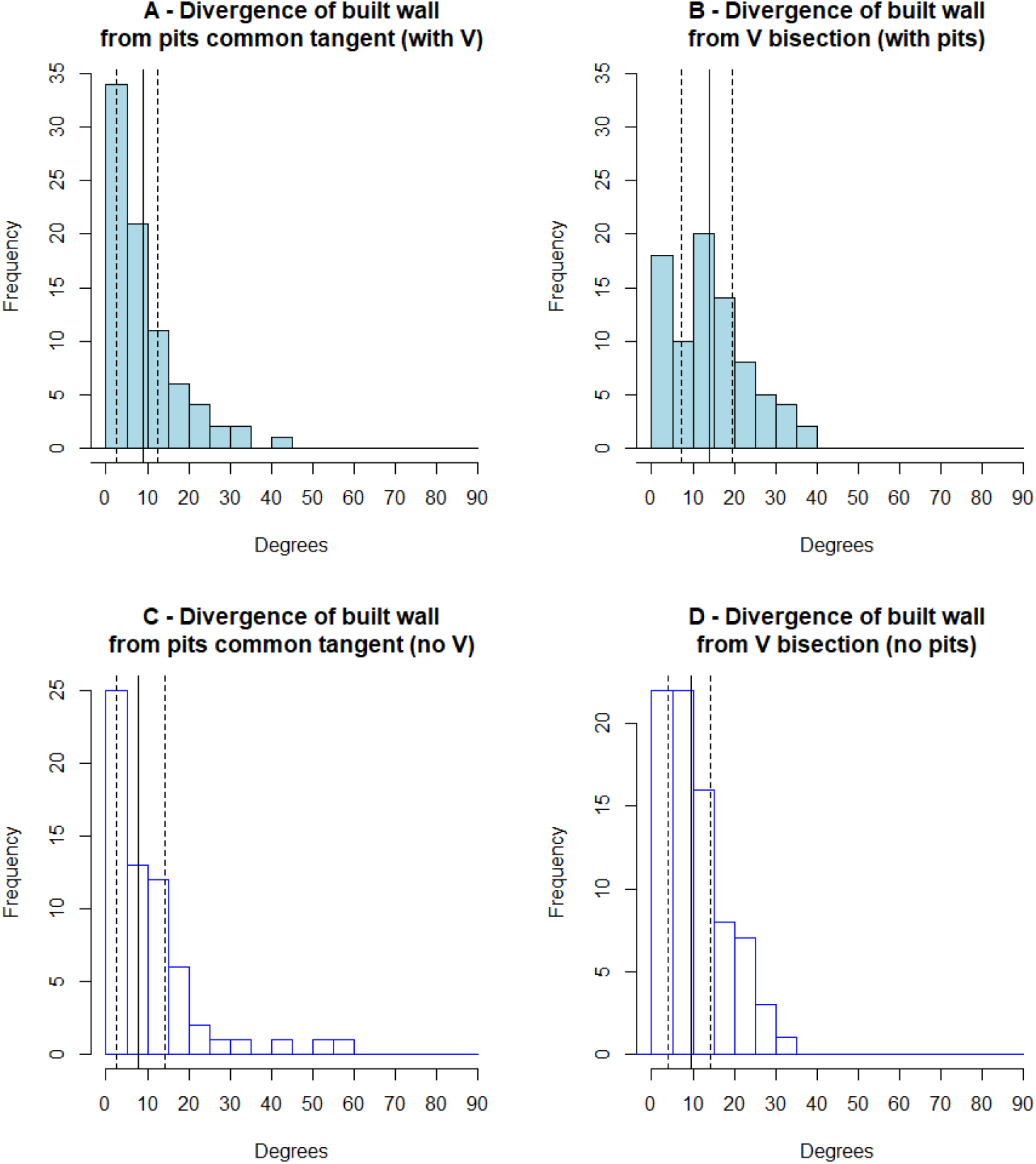
Using compound stimuli comprising both a V-shape and misaligned pair of pits, A shows the distribution of divergence between built walls and the pit pair common tangent while B shows the divergence between built walls and the V stimulus bisection angle. For comparison (copied from experiments 3ii and 3iii) C shows the divergence between built walls and the pit pair common tangent without the additional V-shape and D shows the divergence between built walls and the V stimulus bisection angle without the addition of pit pairs.

Comparing the divergence from the ‘V’ bisection with or without the pits as an additional stimulus, this experiment (14.0°±9.2°, Figure 1-13 B), was significantly different from experiment 2 (10.5°±7.6°, Figure 1-6 A, also Figure 1-13 D ; P = 0.01). This demonstrated that in the presence of the ‘V’ strip, the addition of pits had more influence over cell wall position than may be expected by chance.

## Discussion

Qualitative stigmergy describes how different patterns of behaviour exhibited by an animal, in this case *A. mellifera*, will be triggered as responses when it is confronted by distinct conditions. In this chapter, I have examined the responses and conditions prevalent during the early stages of cell construction. When taken together, the results indicate the presence of a self-organised mechanism that causes initial deposits of wax to be made by builders in a certain manner. The presence of these deposits then induces changes in subsequent builder activity. In this way, builder responses act as stigmergic triggers for future responses, that ultimately contribute to the well-characterised hexagonal structure of comb.

While many publications describe the form of completed cells and the architecture of comb, there are few that address the early stages of construction; a topic to which it is hoped that this chapter contributes. In one of these articles, Huber (1814:139–50) described his observations of the sequence of actions that resulted in the construction of two rows of cells in detail. According to these observations, the first stage of cell construction involved a single worker focusing her efforts on a small depression in a wax deposit, extending it by the removal of wax. Huber also noted that “..the block itself was not of a sufficient length to complete the diameter of the cell. So the bees continued to increase its size” (1814:141). Furthermore, in his description of the beginnings of the second row of cells, Huber observed that the base of a new cell was started by extending the surface formed by a valley at the junction between two extant cells.

In addition to this description of early-stage cell construction, there have been previous attempts to model the steps taken by workers to build comb (Nazzi 2016; Narumi 2018). For example, Nazzi (2016) observed that a cell base would initially be built “in the groove between two pre-existing cells”. The results of experiment 3i support these observations, showing that a depression, or pit, will act as a cue for attention. Stigmergy will then trigger a reaction by the builder to treat such a pit as the beginning of a cell, requiring enlargement.

Once the attention of builders is attracted to a pit, experiments 3i, 3ii and 3iii all show that a wall will likely be built around wax deposited at the edges of the concave stimulus. This empirical outcome supports Huber’s further observations that the initial shallow depressions, while still being enlarged, are worked by other bees that take wax scales “ … and apply them upon the edges so as to lengthen them” (1814:141). Eventually, these edges became cell walls. Nazzi also described the early construction process, stating that “when the cell base is as large as the cell diameter, the walls are started” (2016).

The early stages of cell formation have previously been modelled as an attachment-excavation, where individual actors carve semi-circular cavities within a body of randomly deposited wax leaving a residue similar to natural formations (Narumi 2018). In this model, inter-cell shapes arise through rules that govern the behaviour of the excavators, in contrast to my suggestion involving targeted depositions around a depression, where wax is deposited only where it is needed. Focused placement of wax, rather than bulk deposition and subsequent erosion to form the shape of a cell would seem to require less material and reduced effort, albeit demanding targeted actions that I propose are guided by stigmergy. A point of commonality with Narumi is, however, the erosion mechanism. I include within my hypothesis I that the elemental spherical shape of the cell base is formed by the behaviour described by Martin & Lindauer (1966), involving the envelope prescribed by a bee’s mandibles through the movement of her head, articulated at her neck. This mechanism is also assumed by Narumi to serve as the basis of the shape of excavation.

Predictions P1, P2 and P3 are supported empirically by the content of this chapter, and support my hypotheses concerning the early stages of cellular construction. This is not a complete description of the formation of a cell, nor that of the entire comb, but in subsequent chapters I present further experiments to support the remaining hypothetical reactions to the workpiece and provide a stigmergy-based explanation of cell construction.

